# Circulating bacterial DNA as a tool towards non-invasive biomarkers for colorectal cancer and adenoma

**DOI:** 10.1101/647644

**Authors:** Qian Xiao, Wei Lu, Xiangxing Kong, Yang W. Shao, Yeting Hu, Ao Wang, Hua Bao, Kaihua Liu, Xiaonan Wang, Xue Wu, Shu Zheng, Ying Yuan, Kefeng Ding

## Abstract

**Objective:** The gut microbiota is closely associated with colorectal neoplasia. While most metagenomics studies utilized fecal samples, circulating microbial DNA in colorectal neoplasia patients remained unexplored. This study aimed to characterize microbial DNA in plasma samples and build a machine learning model for colorectal neoplasia early detection.

**Design:** We performed whole genome sequencing of plasma samples from 25 colorectal cancer (CRC) patients, 10 colorectal adenoma (CRA) patients and 22 healthy controls (HC). Microbial DNA was obtained by removing the host genome and relative abundance was measured by mapping reads into microbial genomes. Significant biomarker species were identified in the discovery cohort and built into a random forest model, which was tested in the validation cohort.

**Results:** In the discovery cohort, there were 127 significant species between CRC patients and HC. Based on the random forest model, 28 species were selected from the discovery cohort (AUC=0.944) and yielded an AUC of 1 in the validation cohort. Interestingly, relative abundance of most biomarker species in CRA patients were between CRC patients and HC with a trend towards CRC patients. Furthermore, pathway enrichment analysis also showed similar pattern where CRA patients had intermediate relative abundance of significant pathways compared to CRC patients and HC. Finally, species network analysis revealed that CRC and HC displayed distinct patterns of species association.

**Conclusions:** We demonstrated characteristic alteration of circulating bacterial DNA in colorectal neoplasia patients. The predictive model accurately distinguished CRC and CRA from HC, suggesting the utility of circulating bacterial biomarkers as a non-invasive tool for colorectal neoplasia screening and early diagnosis.

## 1. Introduction

Colorectal cancer is the third most common cancer worldwide with more than 1.3 million new cases and also a leading cause of cancer mortality worldwide[1, 2]. The 5-year survival rate is more than 80% if patients are diagnosed at early stage, but it decreases to about 10% when diagnosed with metastatic colorectal cancer[3]. As early as in 1990, Fearon et al. reported most colorectal cancers were preceded by dysplastic adenomas which would progress into malignant carcinoma, referred to as the classical adenoma-carcinoma sequence[4]. Due to the high incidence of colorectal cancer, the relatively long preclinical phase and identifiable precancerous lesions that could be surgically removed, population-wide colorectal cancer screening programs played a crucial role in reducing overall disease burden and has achieved great success[5]. Currently guaiac fecal occult blood test (gFOBT) and fecal immunochemical test for haemoglobin (FIT) are the standard non-invasive tests for colorectal cancer screening[6]. Other fecal screening tests such as Cologuard which focused on DNA mutation and methylation biomarkers are also emerging[7]. Although these inexpensive and non-invasive tests have good specificity and lead to about 16% reduction in colorectal cancer related mortality[8, 9], they are blamed for comparatively low sensitivity (13% to 50%) for detecting colorectal carcinoma and even more poor sensitivity (9% to 24%) for detecting colorectal adenoma[6, 9]. Hence it is of urgent demand to develop more accurate colorectal cancer screening modalities.

Cumulating studies supported that the progression of colorectal neoplasia was closely associated with gut microbial dysbiosis[10, 11, 12, 13]. Castellarin et al. reported that *Fusobacterium nucleatum* was more abundant in colorectal cancer tissues than normal tissues[10], and it could promote colorectal carcinogenesis by regulating E-cadherin/β-catenin signaling through its FadA adhesion[11]. Furthermore, increasing studies suggested that colorectal lesions harbored a variety of other gut microbiota such as *Bacteroides fragilis, Fusobacterium spp.* and *Roseburia*[14, 15]. Some researchers also made great efforts to establish new colorectal cancer screening models based on the human gut microbiome. Yu et al. identified 20 microbial gene markers in fecal samples that could differentiated colorectal cancer patients and healthy controls, with an area under the receiver operating curve (AUC) of 0.77[12]. Georg et al. built a colorectal cancer early detection model with an AUC of 0.84 by fecal sample metagenomic sequencing, yet they found that fecal microbiota of colorectal adenoma patients was almost indistinguishable from healthy controls[13]. Although these fecal microbiota-based models had moderate sensitivity and specificity for early detection of colorectal cancer, fecal tests usually had poor compliance despite that various beneficial plans had been proposed[16]. Recent liquid biopsy tests such as EpiproColon which detected plasma methylated Septin9 still produced relatively unsatisfactory sensitivity of 0.618[17]. Therefore, non-invasive alternatives for colorectal cancer screening with robust performance and good compliance are urgently demanded.

Recently emerging evidences demonstrated the presence of highly divergent bacterial-specific DNA in human blood, though their source of origin is still a mystery to be solved[18, 19, 20]. Huang et al. reported that blood bacterial biomarkers had prognostic value for early-onset breast cancer patients[21]. Nevertheless, it is not well understood if any correlation exists between blood microbial DNA and disease progressing in other cancers. Given that the gut microbiota has the largest presence of bacteria both in abundance and number of species compared to other organs[22], we hypothesized change in blood microbial DNA composition was associated with colorectal cancer and could be used to identify CRC or HC. Here we presented the pioneer study that utilized whole genome sequencing to characterize circulating microbiome in healthy individuals and patients with colorectal neoplasia. In addition, we constructed a robust classifier model that can accurately distinguish both colorectal carcinoma and adenoma from healthy control.

## 2. Materials and methods

### 2.1. Patients and sample information

A total of 57 participants, including 25 sporadic colorectal cancer patients, 10 colorectal adenoma patients and 22 healthy participants were enrolled in this study. All participants were from the Department of Colorectal Surgery of the Second Affiliated Hospital of Zhejiang University School of Medicine, and the healthy participants have undergone colonoscopies to exclude colorectal neoplasms.

To reduce confounding effects such as age and sex, we enrolled CRC patients and HC with matching demographic features in the discovery cohort. Other CRC and CRA patients treated at the hospital during the study were enrolled in the validation cohort. Specifically, the discovery cohort consisted of 12 CRC patients and 11 matched HC, while the validation cohort consisted of 13 CRC patients, 11 CRA patients and 10 HC (Table 1).

### 2.2. DNA extraction, library preparation and sequencing

Plasma cfDNA was extracted with NucleoSpin Plasma XS kit (Macherey Nagel) using a customized protocol optimized based on the manufacturer’s instructions. Purified DNA was qualified by Nanodrop2000 (Thermo Fisher Scientific) and quantified by Qubit 2.0 using the dsDNA HS Assay Kit (Life Technologies) according to the manufacturer’s recommendations.

Sequencing libraries were prepared with KAPA Hyper Prep kit (KAPA Biosystems) based on the manufacturer’s instructions. In brief, 10ng-200ng of cfDNA were processed by end-repairing, A-tailing and ligation with indexed adapters compatible with the Illumina sequencing platform (Illumina), followed by size selection using AMPure XP beads (Agencourt), PCR amplification with Illumina p5 (5’-AAT GAT ACG GCG ACC ACC GA 3’) and p7 (5’-CAA GCA GAA GAC GGC ATA CGA GAT 3’) primers, and purification by AMPure XP beads.

Quantification of libraries was performed by quantitative polymerase chain reaction (qPCR) using the KAPA Library Quantification kit (KAPA Biosystems). Library fragment size was determined by the Agilent 2100 Bioanalyzer (Agilent Technologies). All sequencing was performed on the Illumina HiSeq4000 NGS platform (Illumina) using paired-end 75bp sequencing chemistry.

### 2.3. Sequence data processing

Trimmomatic was used for FASTQ file quality control (QC). Leading/trailing low quality (quality reading below 15) or N bases were removed. Reads from each sample were first mapped to reference sequence hg19 (Human Genome version 19) using Bowtie2 (v2.3.4.2) with default parameters[23]. Reads that mapped to hg19, mitochondrial genomes or bacterial plasmids were removed (NCBI RefSeq database, accessed on July 19, 2018). The remaining reads were mapped to NCBI microbial reference genome databases using k-mer-based algorithm with Kraken[24]. Relative abundance at bacteria species and genus level were estimated for each sample by Braken with recommended parameters. Results from Braken were fed into HUMAnN2 to generate metabolic pathway level relative abundance for each sample[25].

### 2.4. Statistical analysis

Fisher’s exact test was used to compare categorical variables, while the Student’s t-test and ANOVA were used to compare the distributions of the continuous clinical characteristics across groups. The difference in relative abundance at species, genus, and pathway level between CRC and HC were compared using MaAsLin, which automatically filters sparse data matrix typical of microbial dataset to remove low density species/genus. Confounding factors including age and sex were controlled using MaAsLin parameters for above analysis. Co-occurrence and co-exclusion between species were analyzed with SparCC, which can robustly handle microbiome data compositionality[26]. Species correlations with *p* value less than 0.05 after FDR adjustment were kept. Microbial network plots were generated using iGraph. In addition, clustering of different sample types was visualized via principle component analysis. All statistical analyses were performed in R (v.3.3.2).

### 2.5. Predictive model

To construct a predictive model, random forest recursive feature selection method was used to identify bacterial markers at species level in the discovery cohort. Species with a q value < 0.1 from MaAsLin results were used as input. The model was run under 2-fold cross-validation repeated 50 times. The receiver operating characteristics (ROC) curve and class prediction were also generated by the model. Number of species (feature) selected was plotted against ROC value at each run. The model that generated the max ROC was selected and evaluated in the validation cohort. In addition, the area under the ROC curve (AUC) was used to evaluate model accuracy.

## 3. Results

### 3.1. Characteristics of the participants

The study design was shown in Figure 1 and characteristics of the participants were presented in Table 1. In the discovery cohort, CRP level in CRC patients was below the upper limit of its normal range, implying these patients were not complicated with acute inflammation. Most of the CRC were medium differentiation, and all CRC patients in this study were sporadic CRC with stable mismatch repair proteins expression (pMMR).

**Figure 1.**
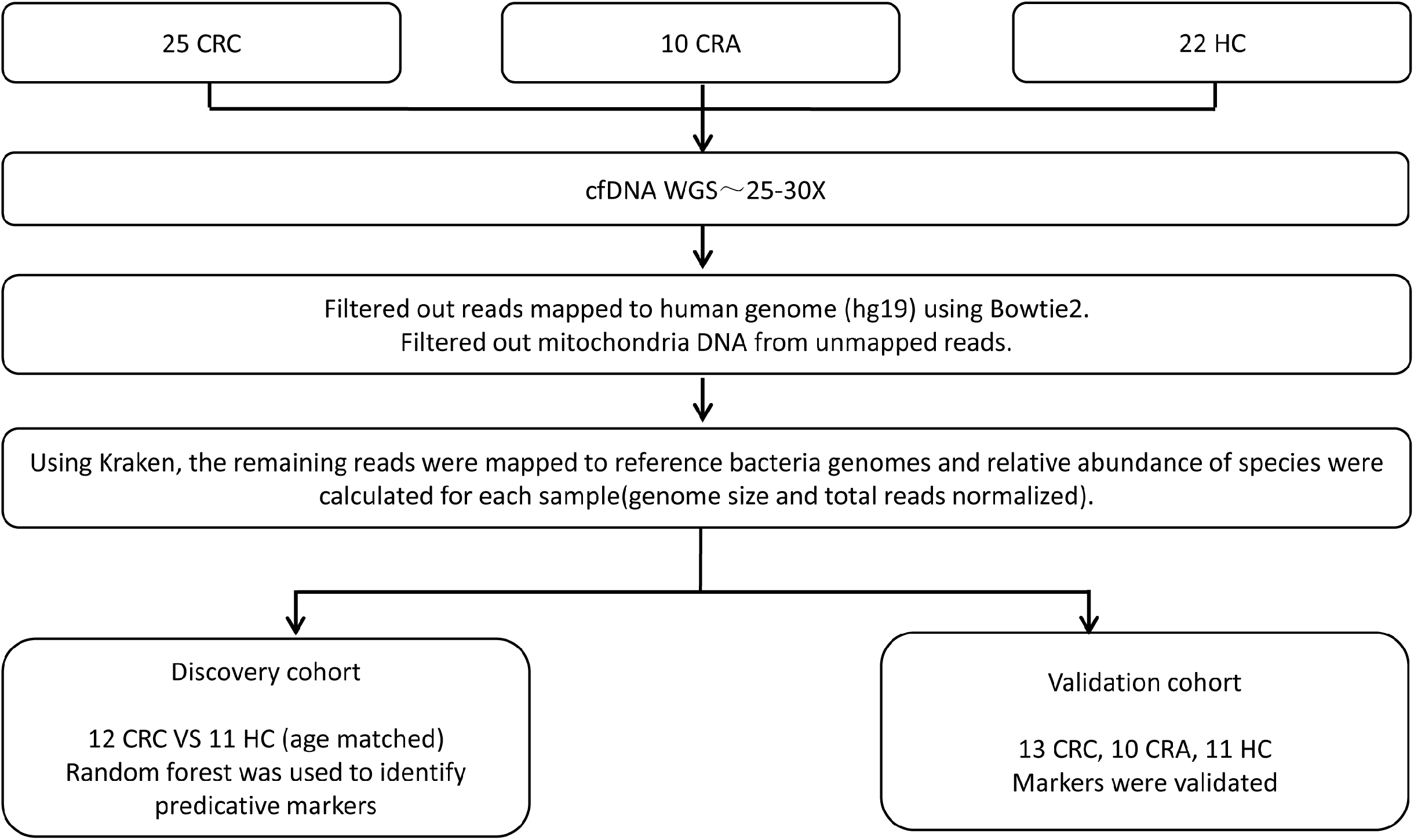
Analytic flowchart. WGS were performed on plasma cfDNA from 25 CRC, 10 CRA, and 22 HC. Reads were filtered to only keep bacterial DNA, which was subsequently mapped to bacterial reference genomes. In the discovery cohort, we identified microbial markers and constructed classifier model to distinguish 12 CRC from 11 HC. In the validation cohort, we evaluated the dichotomous predictive power of the classier model in 13 CRC, 10 CRA, and 11 HC.

In the validation cohort, there was no significant difference among 3 groups in terms of gender, BMI and WBC count, while HC was significantly younger than CRA and CRC patients. There was 5 non-advanced and 5 advanced adenoma in CRA group, and most of the CRC were medium differentiation.

### 3.2. Richness and diversity of circulating microbiome

Species rarefaction curves showed that the number of samples needed to capture the full spectrum of species was slightly different between groups, with CRC and CRA requiring 9 samples while HC requiring 14 samples (Figure 2A-C, Figure S1). There was no significant difference among 3 groups in terms of total number of species, although both shannon diversity and simpson diversity in healthy participants were higher compared to the other two groups (Figure 2D-F). Intriguingly, reduced bacterial diversity in CRC circulating microbiome was consistent with reduced bacterial diversity in CRC fecal microbiome[12].

**Figure 2.**
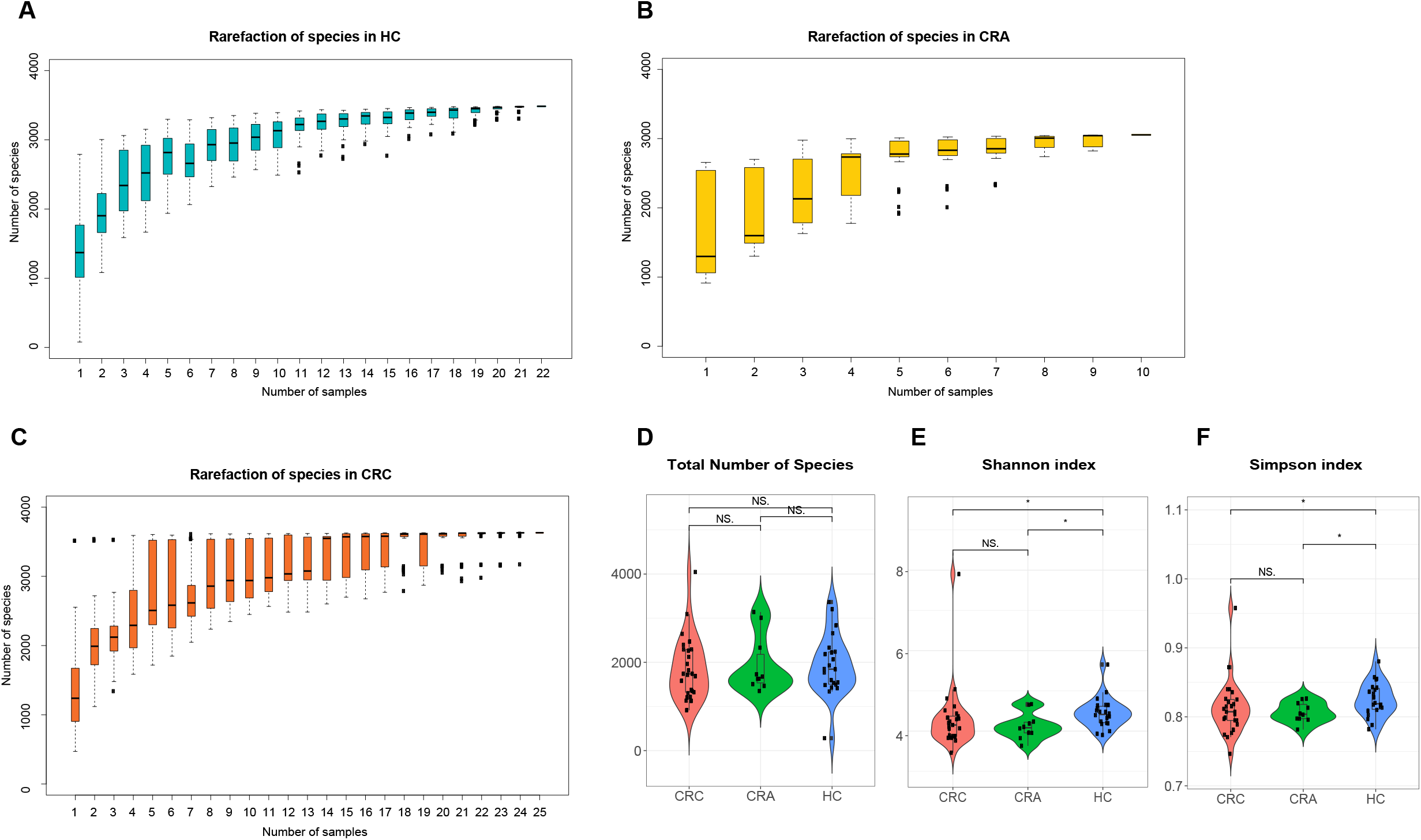
Overview of species distribution at sample and cohort level. Rarefaction curve in (A) HC, (B) CRA and (C) CRC samples. X-axis denoted number of samples that were selected randomly at each run while y-axis denoted the total number of species in the union of samples from that run. Each boxplot represented 50 repetition points. (D) Total number of detected species, (E) Shannon diversity index and (F) Simpson diversity index were computed from all samples in each group.

### 3.3. Abundance profiles of circulating microbiome

Among 3883 identified bacteria species in this study, the majority (3774 species) were present among all 3 groups. CRC, CRA, and HC groups had 20, 1, and 13 unique species respectively (Figure S2). Based on species relative abundance of each sample, principle component analysis (PCA) revealed distinct pattern among 3 groups, where CRA were more close to CRC than HC (Figure 3A). The first and the second principle components explained 11.9% and 8.9% of total variance respectively.

**Figure 3.**
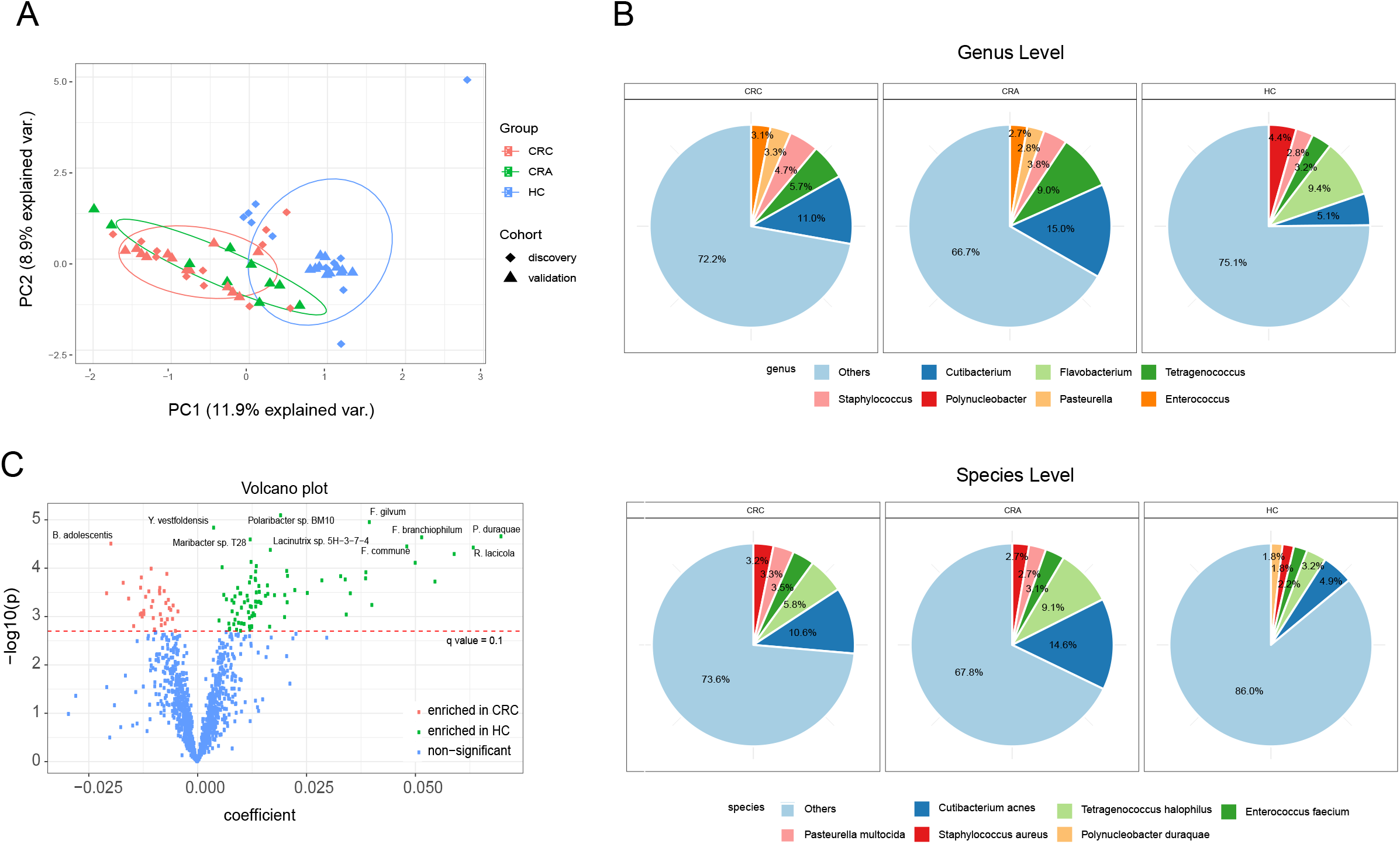
Distinct microbial community composition in CRC, CRA, and HC. (A) Principle component analysis showed stratification of samples by species level relative abundance. PC1 and PC2 values represented the top two principal coordinates. Different sample types were denoted by color code while the discovery and validation cohorts were distinguished by shape. (B) Pie charts showed the most abundance genus (top panel) and species (bottom panel) in CRC, CRA, and HC samples. (C) Volcano plot showed fold change in relative abundance versus −log(p) value for all species in the discovery cohort. Red-dashed line denoted q-value of 0.1. The top 10 most significant species between CRC and HC were labeled.

At genus level, the top 5 most abundant genus accounted for 28% of relative abundance in CRC patients on average, and the proportions were 33% and 25% in CRA patients and HC (Figure 3B). Notably, genus *Flavobacterium*, normally important for mucosa-adherence in gastrointestinal tissues, has been reported to decrease in CRC tissues[27]. Here we also observed a drastic decrease of *Flavobacterium* from being the most abundant genus in HC (9.4%) to less than 1% in both CRC and CRA. We also saw a 10-fold increase of abundance from HC to CRC for genus *Ruminococcus*, which has been reported to be associated with higher CRC risk[28]. At species level, the top 5 most abundant species accounted for 26%, 32% and 24% relative abundance in CRC, CRA and HC respectively (Figure 3B).

In the discovery cohort, differential abundance analysis based on MaAsLin identified 38 species significantly enriched in CRC patients and 89 species significantly enriched in HC (q < 0.1, Figure 3C). The top 5 most significantly enriched species were *Polaribacter sp. BM10, Flavobacterium gilvum, Yoonia vestfoldensis, Polynucleobacter duraquae* and *Flavobacterium branchiophilum*, which were all enriched in HC.

### 3.4. Identifying microbial markers of CRC and independent validation

Based on these 127 significant species, the random-forest model obtained the maximal AUC value of 0.944 when 28 species were selected as features (Figure 4A), half of which were enriched in CRC and half were enriched in HC (Figure 4B). Based on relative abundance of the 28 selected species, PCA revealed more distinct pattern among 3 groups than PCA constructed with total bacterial relative abundance, and CRA largely overlapped with CRC (Figure 4C). The variances explained by principle components were notably improved, with the first and the second principle components explaining 57.7% and 12.3% of total variance respectively.

**Figure 4.**
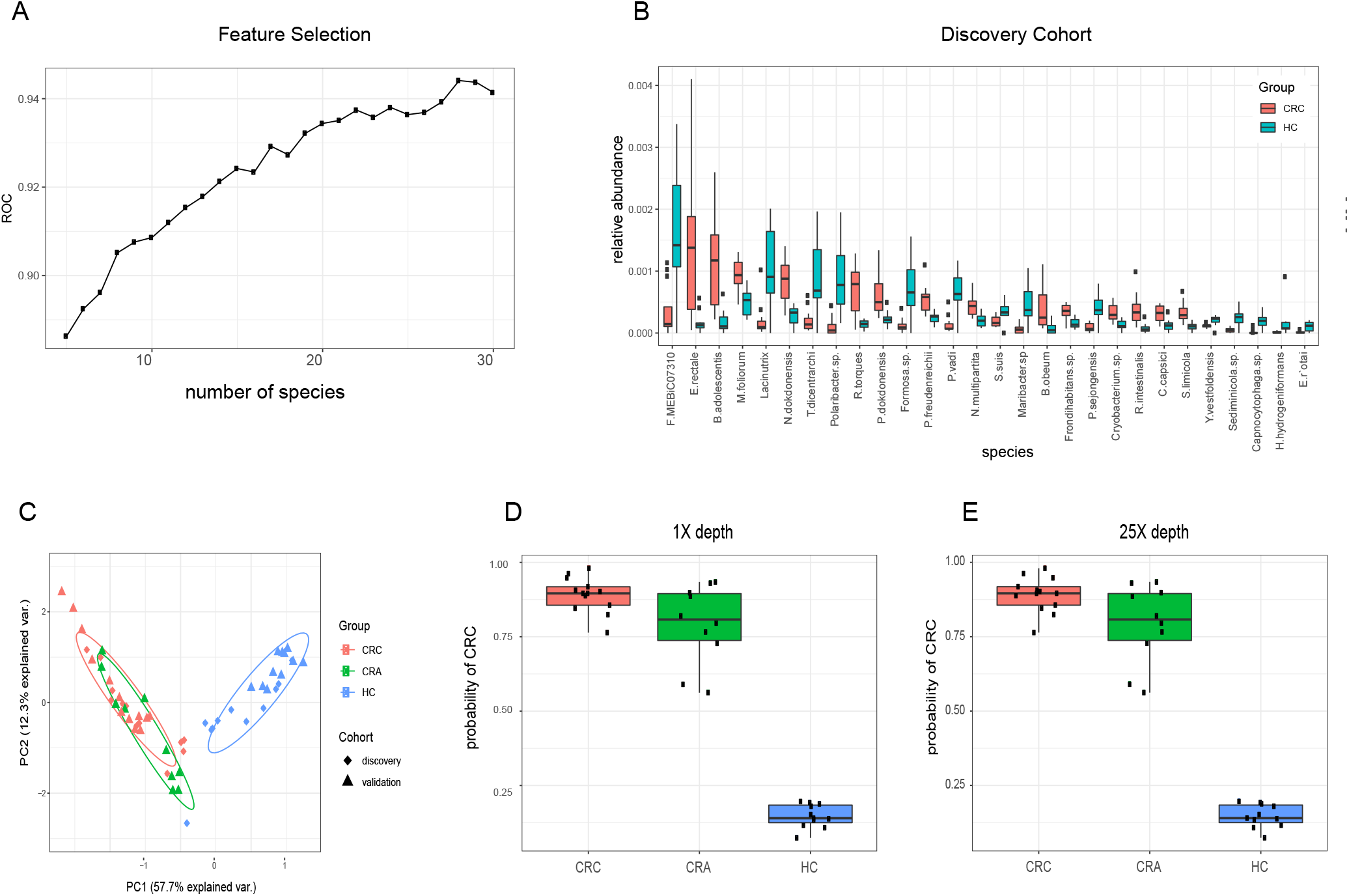
Predictive model construction and validation. (A) In the discovery cohort, species with q-value < 0.1 were used as input to build a classifier model using the random forest recursive selection algorithm. The max ROC value was obtained when the model selected 28 species. (B) Boxplots showed relative abundance of the 28 species in the discovery cohort. (C) Principle component analysis showed stratification of samples by relative abundance of these 28 species. PC1 and PC2 values represent the top two principal coordinates. Different sample types were denoted by color code while the discovery and validation cohorts were distinguished by shape. Model prediction results in the validation cohort were given with (D) randomly down-sampled 1X data or (E) with the original data.

In the validation cohort, this model distinguished CRC patients from HC with an AUC of 1. Importantly, the same model also separated CRA patients from HC with an 100% accuracy. Since the above prediction results were based on the original 30X whole genome sequencing data, we further explored whether the model performance was still robust under the circumstance of low sequencing depth. We first randomly down-sampled the sequencing data to 1X, then analyzed the down-sampled data with the same pipeline as before. Our model still distinguished CRC patients from HC with an AUC of 1, and separated CRA patients from HC with 100% accuracy (Figure 4D-E). Intriguingly, several species abundance in CRA patients such as *Nakamurella multipartite* and *Clavibacter capsici* were intermediate of these in CRC patients and HC, while the majority of the remaining species abundance more closely resembled those in CRC patients than HC (Figure S3). This phenomenon suggested that the classical adenoma-carcinoma sequence was accompanied by the gradual alteration of certain bacteria compositions in the circulating microbiome.

### 3.5. Correlation analysis and pathway enrichment analysis of CRC associated circulating microbiome

We further performed correlation analysis to investigate the relationship among the 28 species in circulating microbiome of CRC patients and HC. We identified 14 species pairs with strong negative correlation (r <= −0.6) and 17 species pairs with strong positive correlation (r >= 0.6) in CRC (Figure 5A and 5C). In HC, 11 species pairs had strong negative correlation and 21 species pairs had strong positive correlation (Figure 5B and 5D). Overall, species enriched in CRC tended to be highly correlated amongst each other and the same trend was also seen in species enriched in HC.

**Figure 5.**
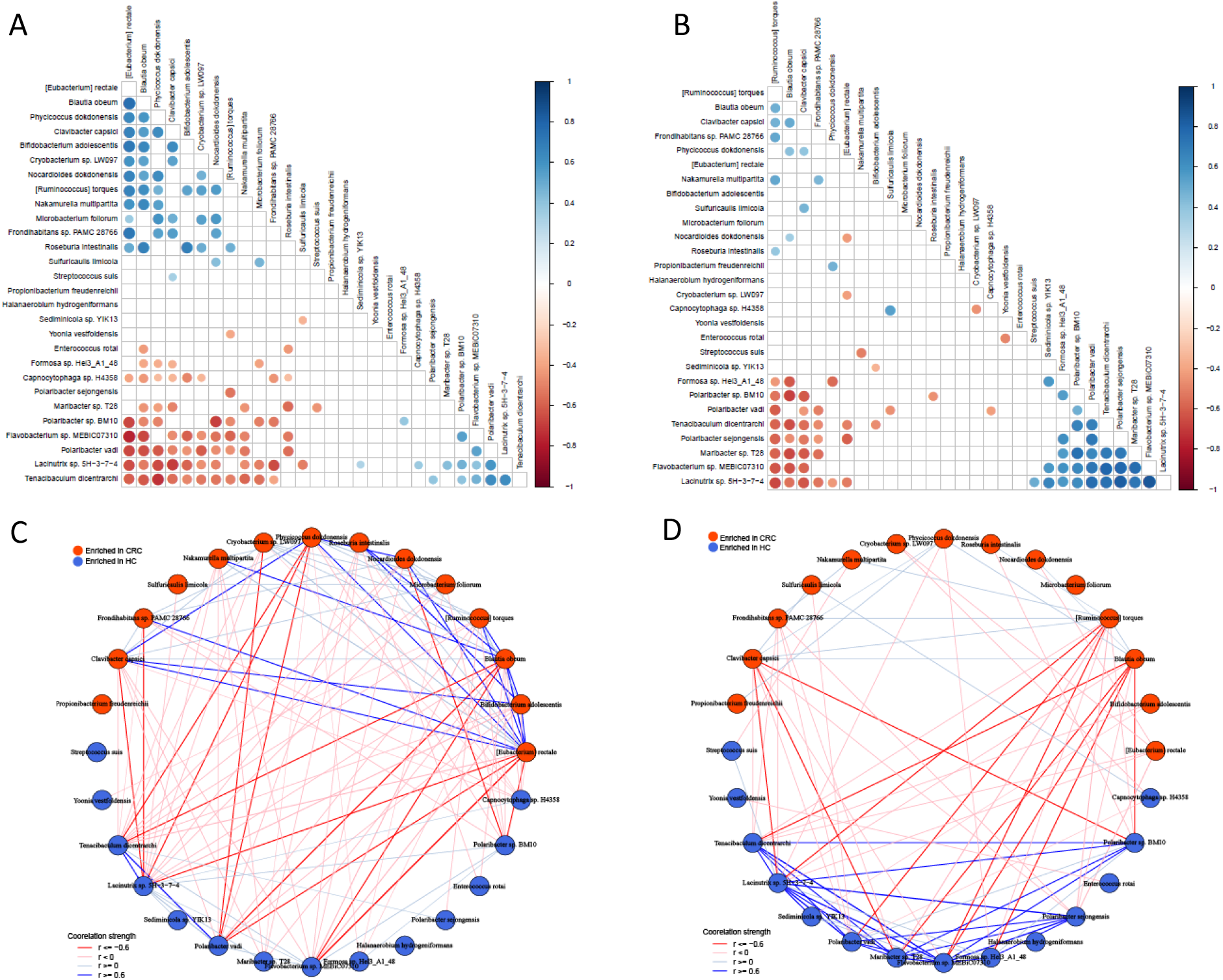
Species correlation in the discovery cohort. Correlation among the 28 species in (A) CRC and (B) HC. Only correlation with *p* value < 0.05 was shown on the plot. The color spectrum represented the strength of correlation. Network plots were constructed to illustrate that (C) CRC and (D) HC had distinct pattern of species level correlations. Red nodes were species enriched in CRC and blue nodes were species enriched in HC. Wider lines demonstrated stronger correlation and only these with *p* value < 0.05 were plotted.

However, we found the pattern of correlation varied a lot in CRC and HC groups. In the CRC patients, the most abundant species *[Eubacterium] rectale* was positively correlated with *Bifidobacterium adolescentis, Blautia obeum, [Ruminococcus] torques, Nocardioides dokdonensis, Phycicoccus dokdonensis, Clavibacter capsici* and *Frondihabitans sp. PAMC 28766*, but it was negatively correlated with *Lacinutrix sp. 5H-3-7-4, Polaribacter vadi, Flavobacterium sp. MEBiC07310* and *Polaribacter sp. BM10.* In the HC group, the most abundant species *Flavobacterium sp. MEBiC07310* was positively correlated with *Maribacter sp. T28, Polaribacter vadi, Lacinutrix sp. 5H-3-7-4, Tenacibaculum dicentrarchi, Polaribacter sp. BM10, Polaribacter sejongensis* and *Formosa sp. Hel3_A1_48*, but it was nagetively correlated with *[Ruminococcus] torques* and *Blautia obeum.* Furthermore, some strong correlation of species abundance in CRC patients, such as *[Eubacterium] rectale* and *Blautia obeum*, were not observed in HC. Furthermore, some correlation of species were completely reversed in these two groups, as *[Eubacterium] rectale* was positively correlated with *Nocardioides dokdonensis* in CRC patients yet negatively correlated with *Nocardioides dokdonensis* in HC. The findings above implied that not only the altered abundance of certain species, but also the dysbiosis of whole bacterial community in the circulating microbiome was possibly associated with CRC pathogenesis.

To investigate whether specific metabolic pathways were altered in CRC, we performed pathway enrichment analysis. MaSaLin identified 20 significant pathways from the discovery cohort (Figure 6A). Notably, TCA cycle pathways were elevated in CRC patients, while petroselinate biosynthesis, superpathway of unsaturated fatty acids biosynthesis (E. coli) and folate transformations I pathways were top enriched pathways in HC. When examined within the validation cohort, these enriched pathways of CRC and HC displayed similar trends as bacterial relative abundance, while CRA patients had values between the other two groups, with closer approximation towards CRC patients (Figure 6B).

**Figure 6.**
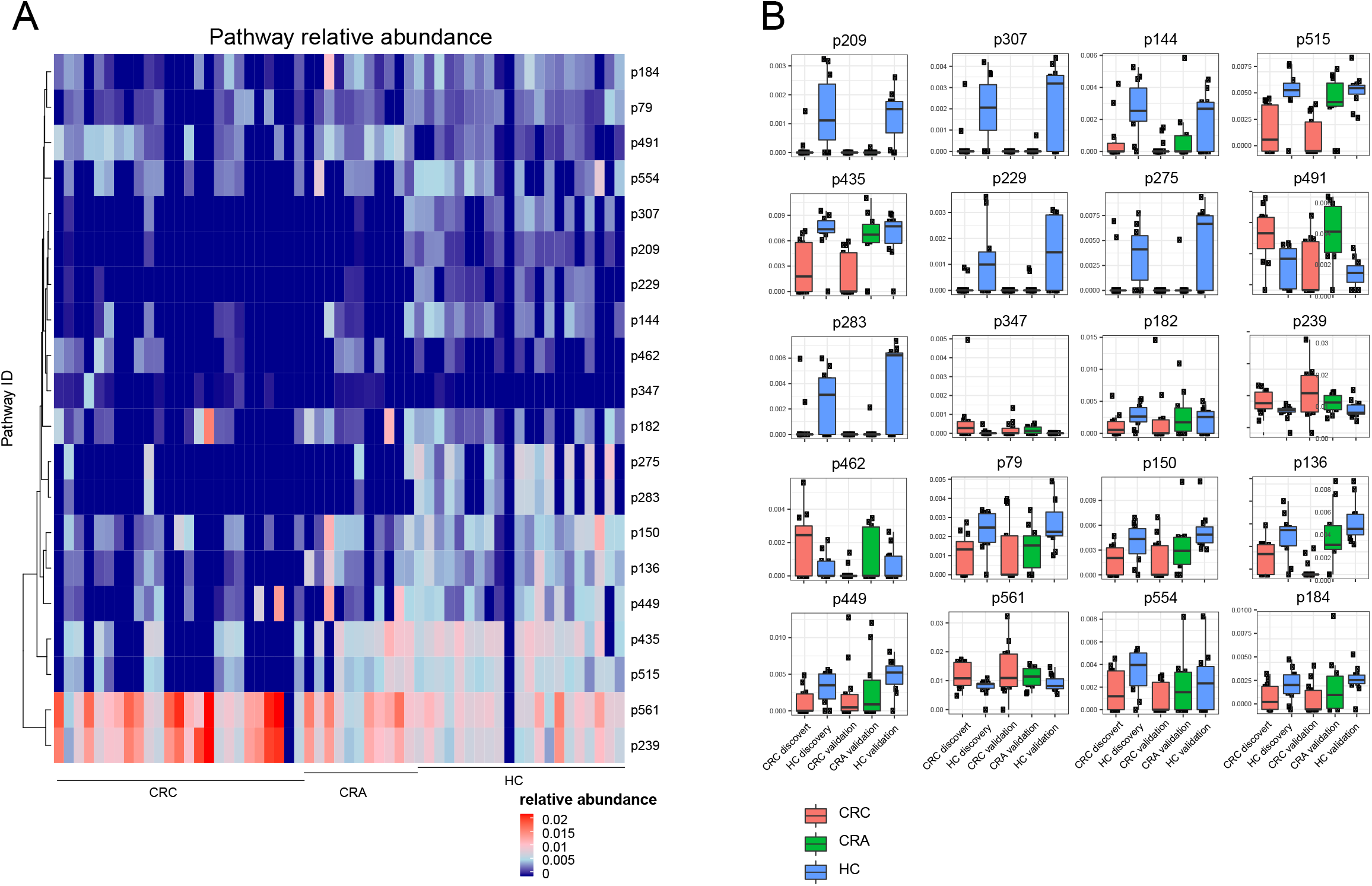
Metabolic pathway abundance. (A) In the discovery cohort, MaAslin identified 20 significant pathways between CRC and HC. We plotted relative abundance of these 20 pathways in all samples with CRC, CRA, and HC clustered for easier comparison. (B) Boxplots showed relative abundance of each pathway grouped by cohort and sample type.

## 4. Discussions

Our study provided the first evidence of distinct blood circulating microbiome composition between healthy individuals and patients with colorectal neoplasia. We found 38 species were significantly enriched in CRC while 89 species were significantly enriched in HC. A robust classifier model based on 28 species could accurately distinguish not only CRC, but also CRA from HC. Species network analysis revealed that CRC and HC displayed distinct patterns of species association. Finally, CRC and HC differed in key metabolic pathways abundance and CRA patients had intermediate relative abundance of these pathways. Altogether, these results indicated drastic change of circulating microbial composition during colorectal tumorigenesis.

Dysbiosis of gut microbiome has long been reported to have close relationship with colorectal cancers, and fecal microbiota-based models had enormous potential to become non-invasive techniques for colorectal cancer early detection and screening[12, 13]. However, circulation as another emerging source of microbiome has recently drawn researchers’ great attention[18], yet the origin of the circulating microbiome is still a mystery. In the current study, we found alteration of several species such as *Flavobacterium* and *[Ruminococcus] torques* were consistent with previously published results in gastrointestinal microbial samples, thus we referred that altered CRC circulating microbiome were partly attributed to dysregulated gastrointestinal microbiome. Nevertheless, a commensal species in oral, *Capnocytophaga sp. H4358*, was also present in the CRC circulation microbiome, indicating that circulation could be a metagenomic pool taking in microbes from other parts of the body.

Recently, tumor liquid biopsy has drawn researchers’ great attention due to a number of advantages including convenience, non-invasiveness and rapid turn-around time. For instance, CancerSeek, which evaluated 8 proteins expression levels and 1933 mutations in diverse genomic positions in the circulation, was designed for the early diagnosis of 8 cancers with an average sensitivity of 62%[29]. In our discovery cohort, 27 species with the highest predictive power were identified by the random forest model, and the classifier model produced accurate prediction of CRC in the validation cohort. More importantly, these predictive markers were able to distinguish CRA from CRC as well, which was difficult using current diagnostic system. Furthermore, robust model performance was produced when sequencing data were down-sampled to 1X from 25X, which implied promising utility of circulating microbial DNA as a diagnostic tool for clinical adoption.

Colorectal cancer was especially suitable for screening due to its long disease course and the remediable precancerous adenoma period, thus researchers have spent a plenty of time to improve the colorectal adenoma screening modality. Currently gFOBT and FIT are the standard non-invasive tests but their sensitivity are less than 50%[6], while other emerging non-invasive tests like aberrant gene methylation detection in fecal samples improve the sensitivity slightly to 63%[30]. Our model demonstrated both high sensitivity and specificity in distinguishing CRA patients from HC even using down-sampled 1X data, which suggested circulating bacterial DNA as a non-invasive and accurate early detection tool for colorectal cancer. In the future, validation of the circulating microbiome-based prediction models in larger cohorts and other ethnics may further improve the stability and efficacy of the current model.

Consistent with the classical adenoma-carcinoma sequence[4], we found several species abundance in CRA patients, such as *Nakamurella multipartite* and *Clavibacter capsici*, were in the middle of CRC patients and HC, while the majority of the remaining species abundance in CRA patients were close to those in CRC patients. Furthermore, significantly enriched pathways in CRA patients were between CRC patients and HC basically. However whether circulating bacterial DNA had molecular functions or were just passengers along CRC carcinogenesis remained unknown.

The current study mainly focused on the circulating bacteria, but did not find significant alteration of other microorganisms such as virus and fungus (data not shown), which were recently reported to be dysregulated in fecal samples of CRC and associated with patients’ overall survival[31, 32]. In addition, despite that CRC was the mostly studied cancer in terms of microbiome, whether dysregulated circulating microbiome was a pan-cancer feature or merely restricted to CRC carcinogenesis was still unexplored.

In conclusion, we demonstrated characteristic alteration of circulating blood bacterial DNA in CRC and CRA patients compared to HC. The predictive model constructed with selected microbial features accurately distinguished CRC and CRA from HC in the validation dataset, suggesting circulating bacterial biomarkers represented a potential non-invasive tool for colorectal neoplasia screening and early diagnosis.

## 5. Acknowledgement

We thank all the kindly volunteers that participated in the study.

## 6. Authors’ contribution

Conception and design: Q. Xiao, W. Lu, X. Kong, Y.W. Shao, K. Ding. Development of methodology: W. Lu, Y.W. Shao, Y. Hu, H. Bao, K. Liu, A. Wang. Acquisition of data: Q. Xiao, K. Liu, A. Wang, X. Wang, X. Wu. Analysis and interpretation of data: Q. Xiao, W. Lu, X. Kong, H. Bao, K. Ding. Writing, review and revision of the manuscript: Q. Xiao, W. Lu, X. Kong, Y.W. Shao, K. Ding. Administrative, technical, or material support: Y. Hu, X. Wang, Y. Yuan, K. Ding. Study supervision: S. Zheng, Y. Yuan, K. Ding.

## 8. Ethics approval

This study was approved by the Ethics Committee of the Second Affiliated Hospital of Zhejiang University School of Medicine (Approval number: 072).

---

**Figure S1. Rarefaction curves for number of species.**

X-axis denoted number of samples that were selected randomly at each run while y-axis denoted the total number of species in the union of samples from that run. Each boxplot represented 50 repetition points. (A) CRC and (B) HC were in the discovery cohort. (C) CRC, (D) HC and (E) CRA were in the validation cohort.

**Figure S2. Venn diagram of number of species detected in three groups.**

**Figure S3. Boxplot of relative abundance of 28 biomarker species in the discovery and validation cohort.**

