## Supplementary figures and images for "Circulating bacterial DNA as a tool towards non-invasive biomarkers for colorectal cancer and adenoma"

### figure S1

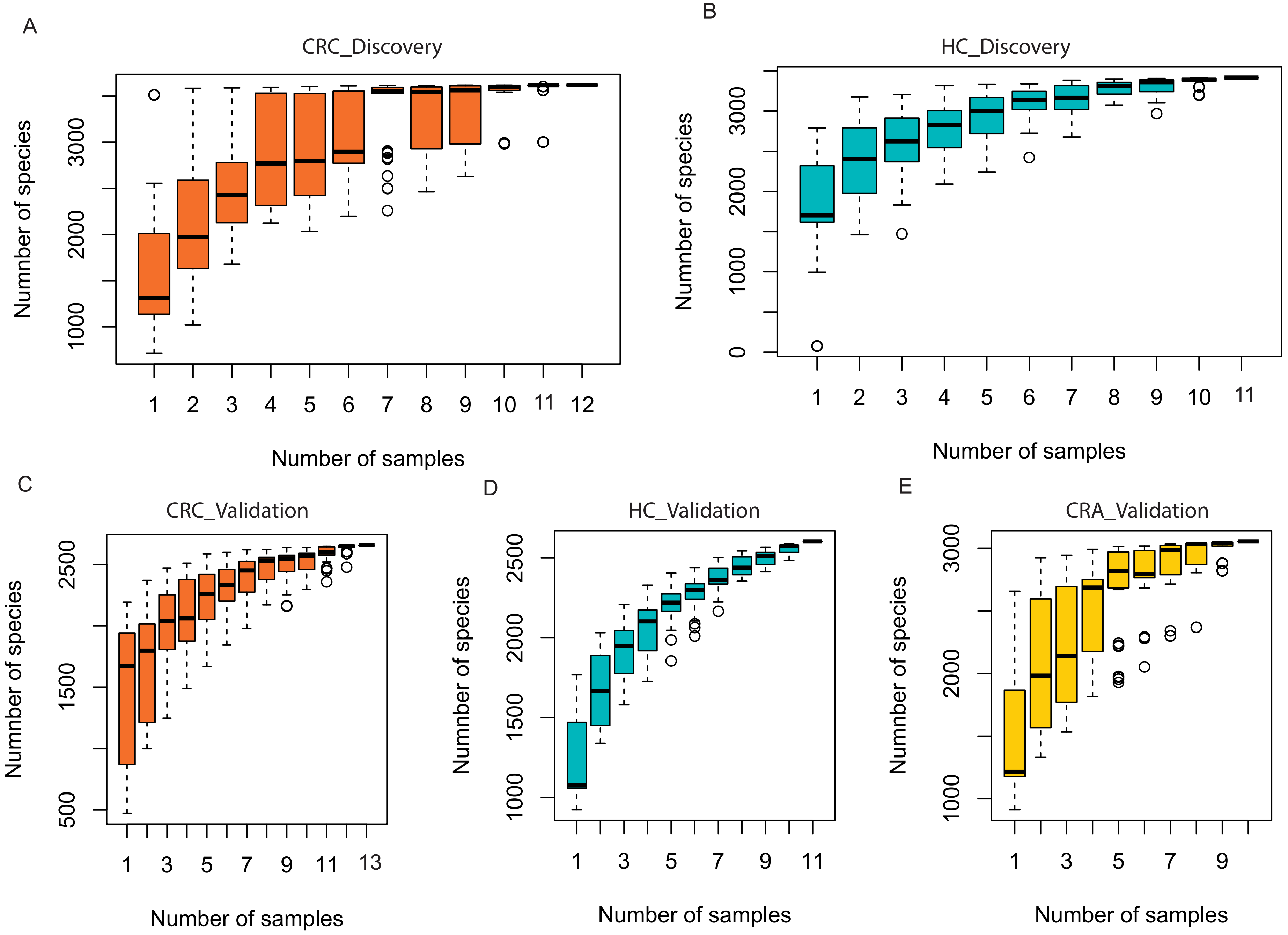

### figure S2

Figure S2

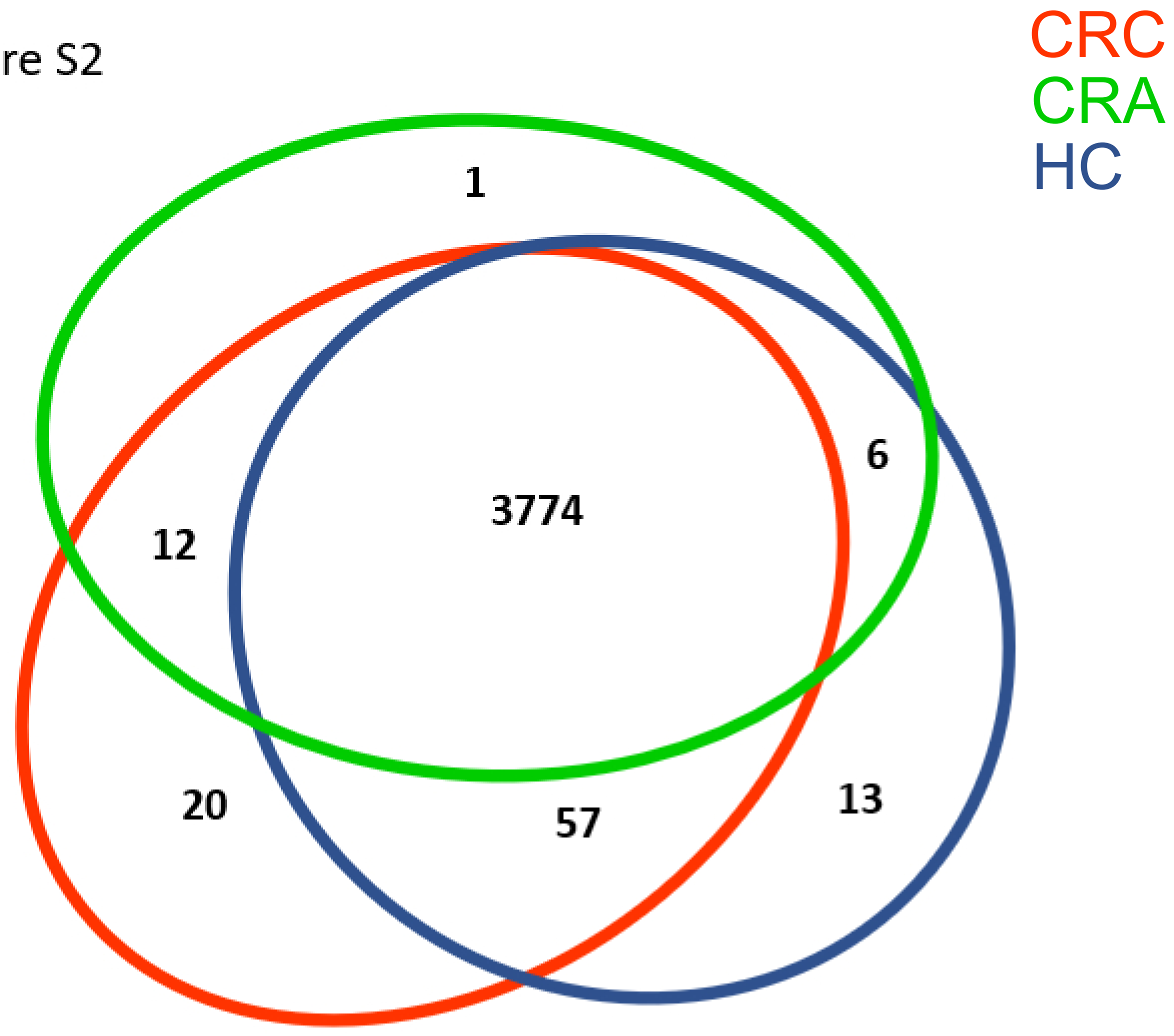

### figure S3

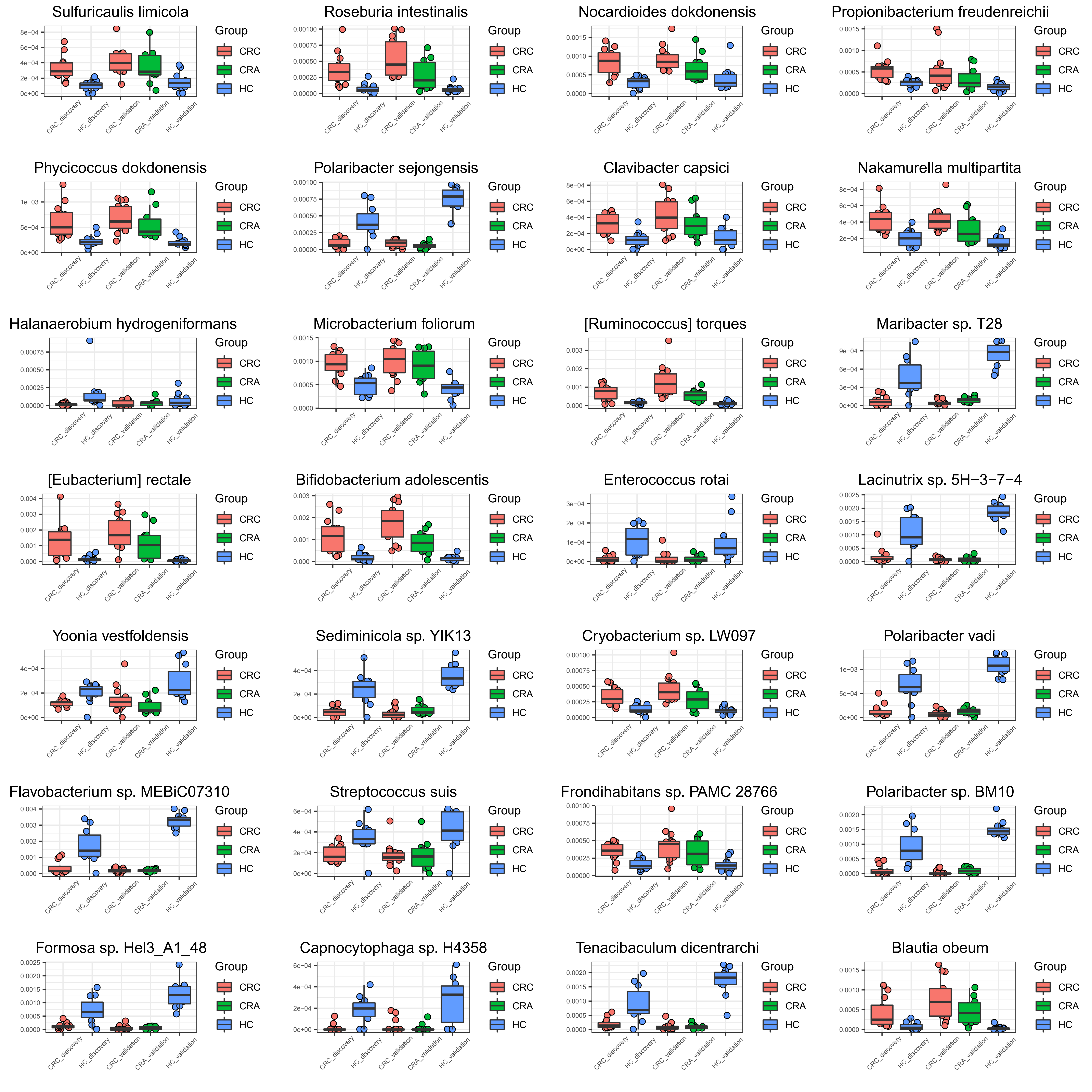
