## Supplementary material for "Circulating bacterial DNA as a tool towards non-invasive biomarkers for colorectal cancer and adenoma": table 1

| Table 1. Characteristics of the included participants | | | | | | | | |
| --- | --- | --- | --- | --- | --- | --- | --- | --- |
| Demographic variables | Characteristics | Discovery cohort | | | Validation cohort | | | |
|  |  | HC | CRC | p | HC | CRA | CRC | p |
| Age (years) | Mean±SD | 49.7±12.5 | 52.2±11.9 | 0.625 | 28.3±1.49 | 62.8±12.3 | 74.6±7.70 | <0.001 |
| Gender | Female | 4 | 4 | 1 | 8 | 3 | 7 | 0.146 |
|  | Male | 7 | 8 |  | 3 | 7 | 6 |  |
| BMI (kg/m2) | Mean±SD | 23.5±3.44 | 23.5±3.43 | 0.985 | 21.2±5.52 | 21.6±2.77 | 20.9±2.86 | 0.907 |
| WBC (*10^9/L) | Mean±SD | 6.18±0.91 | 6.77±2.25 | 0.432 | 5.83±9.17 | 6.46±1.61 | 5.61±1.66 | 0.372 |
| CRP | Mean±SD | NA | 5.17±4.72 |  | NA | NA | 7.78±8.74 |  |
| Fecal OB | Negative | 9 | 2 | 0.003 | 11 | 9 | 0 | <0.001 |
|  | Positive | 2 | 10 |  | 0 | 1 | 13 |  |
| CEA | Mean±SD | 2.00±0.80 | 12.10±27.97 | 0.246 | 1.43±0.32 | 3.53±2.03 | 20.52±22.62 | 0.004 |
| TNM stage | I | NA | 0 |  | NA | NA | 2 |  |
|  | II | NA | 7 |  | NA | NA | 8 |  |
|  | III | NA | 5 |  | NA | NA | 2 |  |
|  | Other | NA | 0 |  | NA | NA | 1* |  |
| Adenoma stage | Non-advanced | NA | NA |  | NA | 5 | NA |  |
|  | Advanced | NA | NA |  | NA | 5 | NA |  |
| Tumor location | Left colon | NA | 4 |  | NA | 2 | 7 |  |
|  | Right colon | NA | 4 |  | NA | 6 | 2 |  |
|  | Rectum | NA | 4 |  | NA | 2 | 3 |  |
|  | Other | NA | 0 |  | NA | 0 | 1* |  |
| Differentiation | Low | NA | 1 |  | NA | NA | 1 |  |
|  | Medium | NA | 9 |  | NA | NA | 11 |  |
|  | High | NA | 2 |  | NA | NA | 1 |  |
| MMR status | dMMR | NA | 0 |  | NA | NA | 0 |  |
|  | pMMR | NA | 12 |  | NA | NA | 13 |  |

*: 1 CRC patient had 2 primary colorectal cancer lesions in the left colon (stage II) and right colon (stage III)
